# FiberAI: A Deep Learning model for automated analysis of nascent DNA Fibers

**DOI:** 10.1101/2020.11.28.397430

**Authors:** Azam Mohsin, Stephen Arnovitz, Aly A Khan, Fotini Gounari

## Abstract

All life forms undergo cell division and are dependent on faithful DNA replication to maintain the stability of their genomes. Both intrinsic and extrinsic factors can stress the replication process and multiple checkpoint mechanisms have evolved to ensure genome stability. Understanding these molecular mechanisms is crucial for preventing and treating genomic instability associated diseases including cancer. DNA replicating fiber fluorography is a powerful technique that directly visualizes the replication process and a cell’s response to replication stress. Analysis of DNA-fiber microscopy images provides quantitative information about replication fitness. However, a bottleneck for high throughput DNA-fiber studies is that quantitative measurements are laborious when performed manually. Here we introduce FiberAI, which uses state-of-the art deep learning frameworks to detect and quantify DNA-fibers in high throughput microscopy images. FiberAI efficiently detects DNA fibers, achieving a bounding box average precision score of 0.91 and a segmentation average precision score of 0.90. We then use FiberAI to measure the integrity of replication checkpoints. FiberAI is publicly available and allows users to view model predicted selections, add their own manual selections, and easily analyze multiple image sets. Thus, FiberAI can help elucidate DNA replication processes by streamlining DNA-fiber analyses.

## Introduction

DNA replication is tightly controlled during the cell cycle to ensure that it occurs only once during S phase and that all the genetic materials are faithfully duplicated, even when conditions are not ideal. Multiple endogenous sources cause replication stress including transcription/replication conflicts, complex DNA structures, and chromatin conformation [1]. Exogenous agents like environmental carcinogens and ionizing irradiation can also cause replication stress and induce genomic instability [2]. To protect the integrity of the genome, checkpoint pathways have evolved that sense replication stress [3]. One important example is the checkpoint-arrest during S-phase which slows down DNA replication to preserve the integrity of the replication fork. During a checkpoint-arrest, cells will attempt to mitigate replication stress including repair of associated DNA breaks before resuming cell cycle progression, and if stress persists cells will undergo apoptosis. Failure of a cell to resolve the replication stress or undergo apoptosis compromises genome integrity through mutations or chromosomal translocations and contributes to malignant transformation [4-6]. Therefore, there is a significant need for robust tools to quantify the integrity and regulation of the DNA replication process.

The DNA-fiber assay is one of the most powerful techniques for the study of DNA replication through the assessment of the integrity of replication checkpoints and the stability of the replication fork. The assay involves incorporation of nucleotide analogs like iododeoxyuridine (IdU) and chlorodeoxyuridine (CldU) into the nascent replicating DNA. Incorporation of the nucleotide analogs to the replicating DNA is then visualized by fluorescent microscopy and the lengths of the nascent DNA-fiber are quantitively measured [7]. Replication stress inducing agents like hydroxyurea, which blocks DNA synthesis by preventing expansion of the dNTP pool, can be applied to cells in various combinations during nucleotide analog incorporation to test the effectiveness of checkpoint stimulation or the protection of the replication fork from degradation [8]. Taken together, the DNA fiber assay is a powerful tool that can be used to directly interrogate the replication machinery in many contexts including cell cycle checkpoints, DNA damage responses, replication fork protection, and response to exogenous agents and therapeutics.

Measuring hundreds of DNA fibers within microscopy imaging can be done manually with general image-processing tools like ImageJ, but this can be extremely tedious and subject to experimental bias. Furthermore, fairly significant number of fibers per condition must be counted to be certain it is reflective of real biology and representative of the population over individual cells. Purely automated methods for fiber analysis, such as through FiberQ, have demonstrated low correlation between automated and manual fiber length estimates and have been found to select 27% more ‘ambiguous’ or extra fibers than manual annotators [9]. Thus, there is a significant need for an accurate and efficient fiber analysis tool. We suggest a middle ground where a tool automatically performs fiber analysis, but also allows for manual intervention to enable experimenters to visualize automated fiber selections, remove any selections they deem incorrect, and allow for seamless addition of custom fiber annotations.

Recent progress in deep learning has made it more efficient to detect subtle patterns in complex images and promises to help automate fiber detection and quantification. In particular, we formulate two image analysis tasks in the context of fiber analysis: object detection and semantic segmentation (Figure 1). First, object detection is used to select and draw a 4-sided polygon, termed bounding box, around every fiber. Second, semantic segmentation, which classifies every pixel to a specific category [10]. In the context of the fiber objects, semantic segmentation is used to classify each pixel in the bounding box as “fiber” or “non-fiber”. In this case, all pixels belonging to the fiber class will be highlighted and enable measurement of fiber lengths. The instance segmentation task is a combination of the object detection and semantic segmentation tasks. It enables the detection of each object with a bounding box and the classification of each pixel inside the predicted bounding box [11]. Thus, instance segmentation defines the framework for automated detection and measurement of fibers from microscopy images, since fiber measurement requires detection of a fiber and the pixels for each fiber.

**Figure 1:**
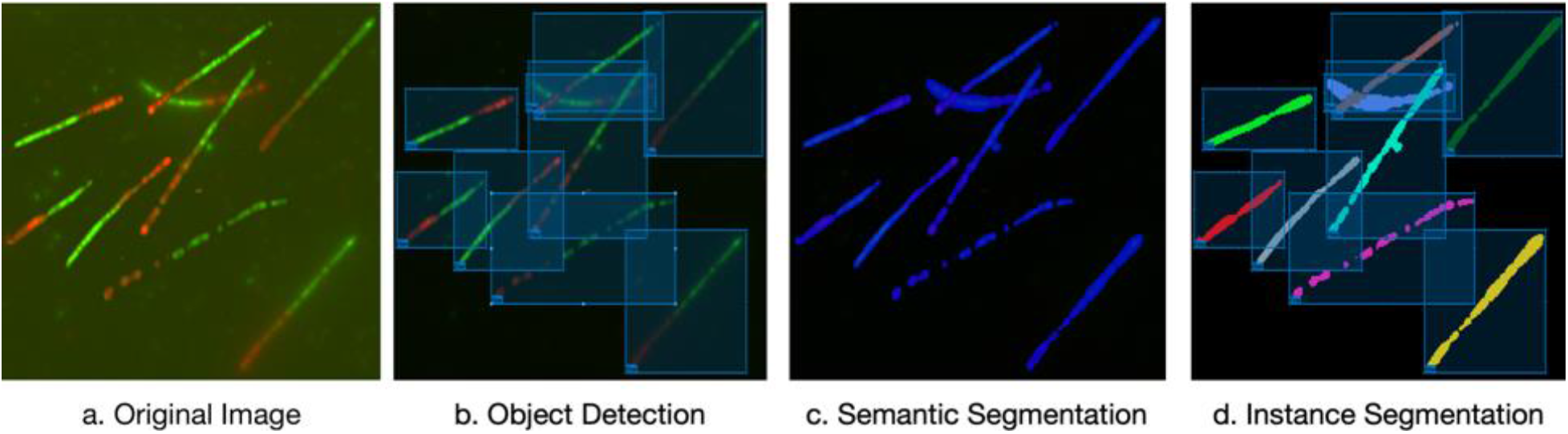
Three relevant computer vision tasks in the context of fiber analysis: Panel A shows the original RGB image with multiple overlapping fibers. Panel B shows predictions from the object detection task where the coordinates of 4-sided bounding box is predicted for each fiber object. Panel C shows the predictions from the semantic segmentation task where each pixel is classified into the fiber or non-fiber category. Pixels in classified into the fiber category are highlighted blue while pixels classified into the non-fiber are highlighted black. Panel D shows the predictions from the instance segmentation task. In instance segmentation (a combination of object detection and semantic segmentation), the coordinates of a bounding box are predicted for each fiber object and then the pixels inside each bounding box are classified into a category. This allows for detection of multiple fiber instances and the pixels belonging to each different fiber instance.

Here we present FiberAI, a platform that includes an instance segmentation model that provides automated selections for two-component DNA fibers and a graphical interface for user visualization and input. For training, we use our own fiber microscopy images and publicly available images. Our trained model is available to run separately in a Python script on a set of images while the graphical interface is created as a MATLAB app. Our software is freely available and all associated code and data can be found at https://github.com/awezmm/FiberAI. Finally, we validate FiberAI by using it to efficiently analyze specific fiber assays that measure replication checkpoint integrity in T cells.

## MATERIAL AND METHODS

### Nascent Fiber Assays

Thymocytes for the Nascent Fiber Assays were isolated from 6-8 week old mice in 2%FBS in PBS and dissociated to single cells using 0.45μM strainers. Cells were then resuspended (IMDM, 10%FBS, 1%Pen/Strep, 50mM β-mercatopethanol) and acclimated for 1 hour at 5% CO2 and 37°C. For untreated fibers, cells were pulse labeled first with 25 μM IdU (MP Biochemicals, 0210025701) for 25 minutes and then with 250 μM CldU (Sigma, C6892) for an additional 25 minutes. For fork recovery assays, cells were cultured in 2mM hydroxyurea (Sigma, H8627) for 3 hours between IdU and CldU pulses. For stressed replication assays, cells were co-cultured in 250uM CldU and 2mM hydroxyurea during the second pulse. After labeling, cells were immediately collected on ice and diluted to 7.5 x105 cells/mL. Cells were then spotted onto the top of glass slides, briefly dried, and lysed by adding buffer (100mM Tris pH 7.5, 0.5% SDS, 50mM EDTA) dropwise. After 2-3 minutes, fibers were spread by tilting slides at a 20-40° angle allowing droplets to roll down the slide length. Slides were then air dried for 30-60 minutes, fixed in a 3:1 methanol/ acetic acid solution for 10 minutes, and denatured in 2.5M HCl for 80 minutes. Fibers were stained at 4°C overnight in a humidified chamber with rat-α-BrdU (1:200, BD Biosciences, B44) and mouse-α-BrdU (1:25, Abcam, BU1/75(ICR1)) in 1%BSA, which label IdU and CldU, respectively. Cells were then washed in PBS, fixed for 10 minutes (3% PFA, 3.4% sucrose in PBS), and stained with α-rat Alexa Fluor 488 (1:500, Invitrogen, A-11006) and α-mouse Alexa Fluor 594 (1:400, Invitrogen, A-11005) for 1.5 hours at room temperature. Slides were then washed and mounted with ProLong Anti-fade mounting media (Invitrogen, P36930) and #1.5 cover glass. Images were acquired with an Olympus IX81 inverted microscope with a 100X, NA 1.45 objective at the Integrated Light Microscopy Core at the University of Chicago.

### Dataset Preparation

To include a variety of fiber staining patterns, our dataset includes images taken from four different DNA fiber assay samples and images from the public repository associated with FiberQ [9]. The raw grayscale image channels from the DNA Fiber assay were merged into RGB images. Images from FiberQ’s public repository were already provided as merged RGB images. All images were resized to height and width dimensions of 1024 and 1024 pixels. The contrast of the red and green channels in the RGB images was increased by mapping the intensity values in each channel to new values by saturating the bottom 1% and top 1% of pixel values using the imadjust function in MATLAB. Fibers were manually labeled with LabelMe, which allowed for easy labelling of the bounding box coordinates of fibers and labelling of the pixels inside the bounding box, belonging to the fiber object [12]. To establish background, pixels with a value less than 0.95 standard deviations above the mean of the Value channel in the HSV (Hue, Saturation, Value) colorspace, were set to a (0,0,0) intensity value in the RGB colorspace.

After manual labelling with LabelMe, we generated a dataset with the adjusted RGB images and with a json file that contained the coordinates of the bounding boxes and pixels for each labeled fiber in each RGB image. At this point, our dataset included 114 manually segmented fibers. Next, the dataset was partitioned into 2 sets: a training set and a test. 85% of the original dataset was partitioned into the training set and the other 15% was partitioned into a test set. In order to increase the size and complexity of the dataset, CloDSA, a Python package that can apply data augmentation to an instance segmentation dataset [13], was used to apply the following image augmentation techniques on the training set: flipping in the horizontal and vertical directions, rotation by 90, 180, and 270 degrees, random shearing, and random translation in the x and y directions. After data augmentation, the final training set included 684 manually labelled fibers across 180 images.

### Model Training and Evaluation

After the dataset preparation, we trained an instance segmentation model to predict the bounding boxes and segmentation pixels for fiber objects. The model was trained Detectron2 which is a next generation software system, powered by PyTorch, an open-source machine learning library [14], that implements state-of-the-art object detection algorithms [15]. First, Detectron2 was used to initiate a pretrained Mask RCNN R 50 FPN model, which is a baseline model previously trained with Detectron2, that has been show to provide one of the best tradeoffs between speed and accuracy [15, 16]. Our model training parameters included a batch size of 2, the base learning rate was 0.02 and was decreased by 0.01 every 700 iterations, and a stochastic gradient descent optimizer. The model was trained for 10,000 iterations on a NVIDIA Tesla T4 GPU, which helped accelerate the rate of preprocessing and training. After training, the model was evaluated for average precision using the COCOEvaluator class available in Detectron2, which calculates average precision using the methods provided by COCO [17]. COCO is a large-scale object detection and segmentation dataset that provides methods for evaluating instance segmentation models.

The ground truth is established through the manual annotation of all images. Precision is defined as: True Positive / (True Positive + False Positive). In the context of the fiber bounding box task, a True Positive prediction is when the model predicts a bounding box around an object that was actually a fiber while a False Positive prediction is when the model predicts a bounding box around an object that was not a fiber. For the bounding box task, Intersection over Union (IOU) can be defined as the area of overlap between the actual and predicted bounding box divided by the union between the actual and predicted bounding box [17]. COCO suggests different IOU thresholds to determine if a bounding box is a True Positive or False Positive [17]. For instance, an IOU threshold of 0.5 would be mean that a bounding box would only be considered a True Positive if it’s IOU against the actual bounding box was at least 0.5. Here, we used Detectron2’s evaluator module to give several precision metrics.

### Measurement Pipeline

The trained model provided bounding box coordinates for each fiber and predicted the pixels that were part of the fiber in each bounding box. A pipeline was then created in Python to measure the red and green components for each fiber bounding box. For each bounding box and segmentation mask, the entire input image was cropped to the coordinates of the bounding box (Figure 2a, 2b). The model also provided a binary image that indicated which pixels were part of the fiber object. The binary object containing the pixels of the fiber was first thinned by using the *skeletonize* function in the scikit-image Python library [18] (Figure 2c). Thinning a binary object refers to reducing the object to a line that has a width of 1 pixel. The color of each pixel in the skeletonized binary object was either changed to a red color (RGB=1,0,0) if its intensity value in the red channel was greater than or equal to its intensity value in the green channel or a green color (RGB=0,1,0). Pixels that were black (RGB=0,0,0) remained black. This created a skeletonized fiber with multiple red, green, or black segments where black segments represented small gaps in the fiber.

**Figure 2:**
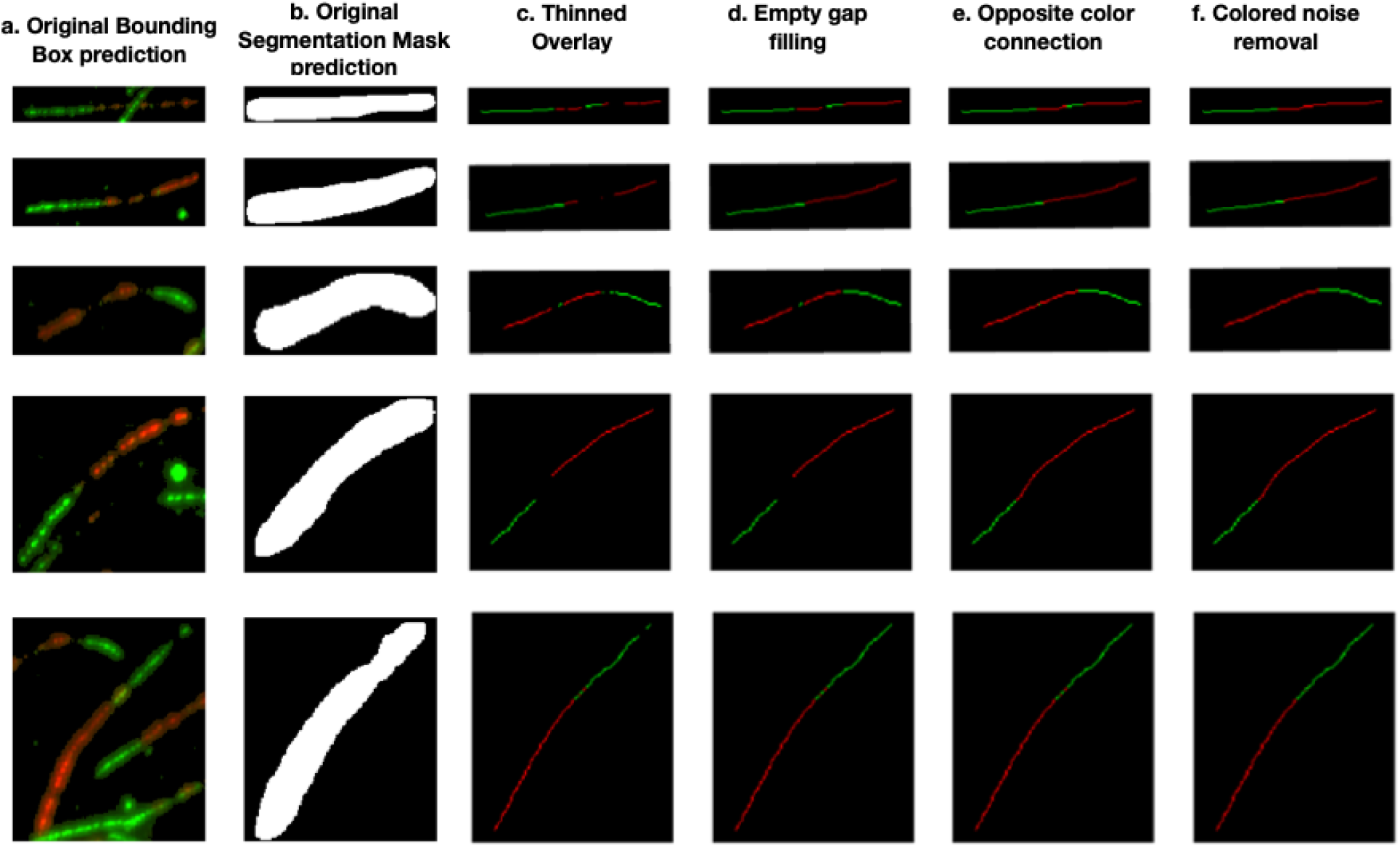
Measurement Pipeline. After bounding box and segmentation, a measurement pipeline involving 4 steps find the lengths of the red and green components of a fiber. Panels A and B show the original bounding box and segmentation mask predictions. The first step of the measurement is to thin (Panel C) which reduces the binary object to a single line of 1-pixle width. Next, any black gaps between 2 larger segments of the same color are replaced with that same color (Panel D). Row 1, Panel D shows an example of a gap near the end of the fiber, between 2 red segments being filled in with red. Any remaining black segments are replaced by the color of the larger segment on either side of it (Panel E). Finally, any segments with a color different than both segments around it and smaller than both segments around it are replaced with the color of the segments around it (Panel F). Row 5, Panel f shows small red segments in the middle of the large green segment being replaced by green. After this processing pipeline, the number of red pixels and green pixels is simply counted to determine the length of the red and green components.

Since, we were measuring two-component fibers only, the skeletonized fiber image needed to be corrected to include only a single continuous red segment and a single continuous green segment, without any black gap components. First, any black components that were between two segments of the same color and were smaller than those two segments were changed to the color of those two segments (Figure 2d). Second, any remaining black segments were changed to the color of the longer of the two segments around it (Figure 2e). This left a fiber without any black segments and multiple red and green segments. Finally, any colored segment that had different color than both segments around it and was smaller than both segments around it was changed to the color of the segments around it (Figure 2f). This ensured that the final fiber had a single continuous red segment and a single continuous green segment.

## Results and Discussion

### FiberAI selects fibers with high precision and accuracy

The FiberAI pipeline includes an instance segmentation model that identifies the pixels that are part of each fiber instance, and subsequently classifies and measures the lengths of the components in two-component fibers. We evaluated the accuracy of the instance segmentation model and the automated measurement separately. The standard method to evaluate an instance segmentation model is to calculate precision values for a bounding box prediction and the pixel segmentation on a test set, as indicated by COCO’s standard evaluation metrics [17]. As explained in methods, the ground truth was established through the manual annotation of all images. We provide precision metric scores out of 1 for the bounding box task and segmenation task(Figure 3a). Here, average precision represents the average of all precision scores over different overlapping proportions (referred to as Intersection Over Union (IOU)) threshold values from 0.5 to 0.95 (step size 0.05). The FiberAI model shows high performance on both the bounding box and segmentation (Figure3a), as the AP for the bounding box and segmentation are 0.91 and 0.90 respectively.

**Figure 3:**
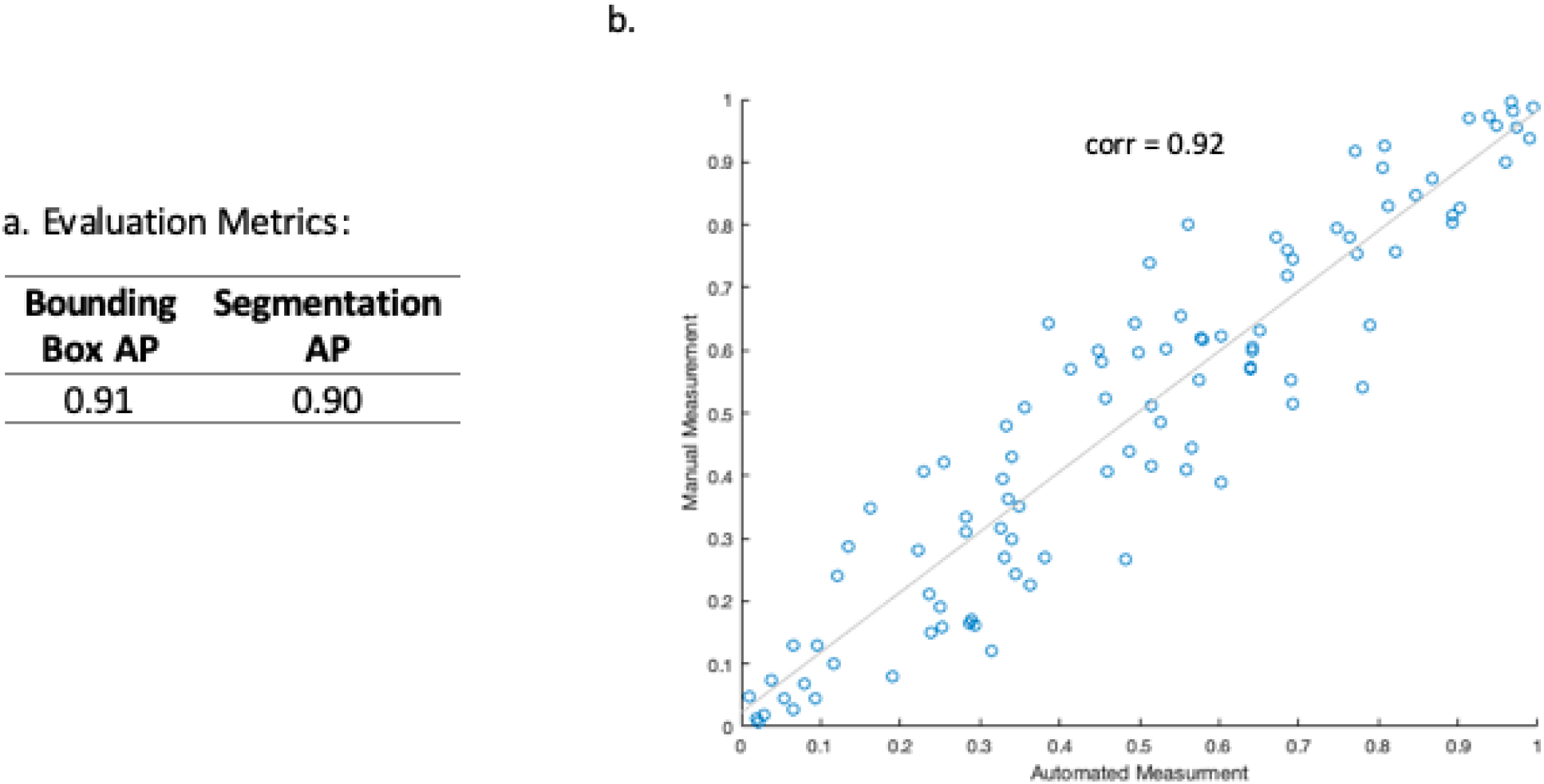
FiberAI evaluation metrics: Panel A shows mean precision values for the bounding box and segmentation tasks. AP refers to mean of all average precision values at IOU thresholds from 0.50 to 0.95 (with a step size of 0.05). IOU refers to the Intersection of Union score between the actual and predicted object. Calculation for precision was conducted using the CocoEvaluator in Detectron2’s evaluation package: https://detectron2.readthedocs.io/modules/evaluation.html. Panel B shows high correlation (0.92) between automated and manual measurements of DNA fibers.

**Figure 4:**
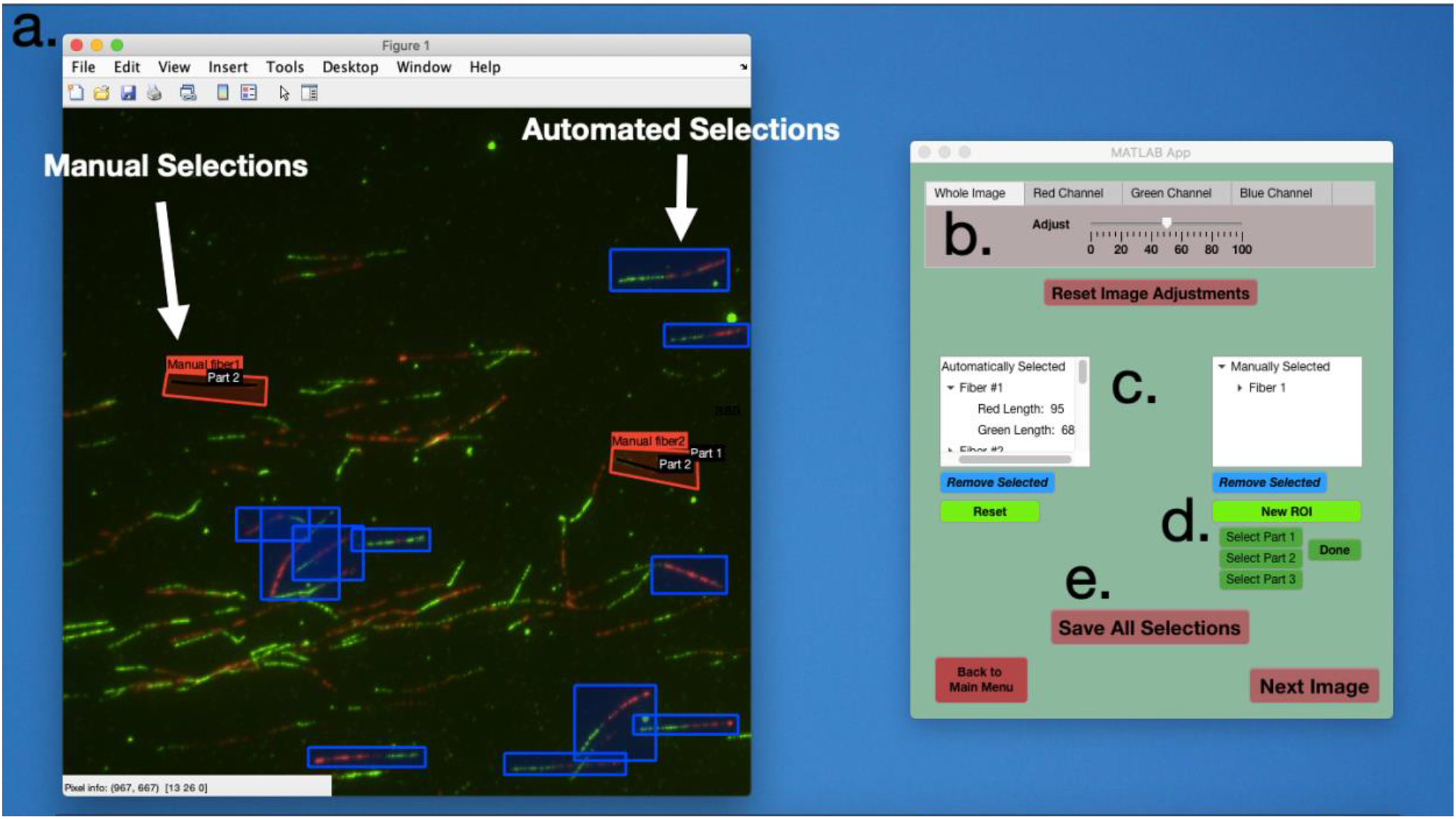
FiberAI Graphical Interface: The FiberAI graphical interface displays each image along with predicted fibers. It also allows a user to remove automated selections or add their own manual selections. Area A shows the original image display with the automated selections (in blue boxes) and manual selections (in red boxes). The toolbar on top of the image allows for zoom controls. A separate window on right contains controls for adjusting image contrast and removing automated selections and adding manual selections. Area B shows the control panel for adjusting the contrast in red, green, blue channels or the whole image. Area C shows two text lists that display automated selections (on the left) and manual selections (on the right). Each item in the text list represent one fiber and each item can be clicked to show the measurements of the components of each fiber. Items can also be deleted by selecting the item and clicking the ‘Remove Selected’ button. Area D shows controls for adding a manual selection. Manual selections can include up to 3 components. Area E shows the control for save all current fiber selections for the current image.

We next evaluated the accuracy of fiber lengths in the automated selections by comparison to measurements performed manually. Here we show that FiberAI-determined measurements from the post-processing pipeline are highly correlated to human performance (Figure 3b, r = 0.92).

### FiberAI graphical interface allows for easy fiber selection manipulation and visualization

The FiberAI graphical interface is built with the MATLAB AppDesigner and can be installed on a Windows or OSX operating system with MATLAB 2020a runtime driver. Users first run a Python script that will use the trained model to predict fiber selections on a set of images and output a .mat file that holds metadata information for the automated selections. Next users can open the graphical interface, load images, and the metadata .mat file. For each image, FiberAI displays the image (4a) and draws predicted selections. Four sliders allow the user to change the contrast of the red, green, blue channels, or of the whole image (4b). Two dropdown lists display automated and manual fiber selections (4c). A user can click on a fiber selection in a list, and use the delete button to remove the selection from the displayed image and the list of automatic or manual selections. Users can manually select a new fiber by clicking the ‘New ROI’ button, drawing a rectangular section on the image, and then clicking on at least one ‘new part’ button to draw line on the fiber to measure a specific section of the fiber (4d). Finally, users can save all selections which will output fiber measurement into an excel file and save an image with region of interest overlays for automatic and manual selections (4e). Users have the ability to select and IOU threshold for the model prediction.

### Using FiberAI to evaluate the integrity of the replication checkpoint

FiberAI was used to measure DNA fibers from an untreated sample and a sample that had been treated with hydroxyurea. The hydroxyurea treated sample serves as a positive control to ensure that FIberAI faithfully detects replication stress resulting in a decrease in the IdU: CldU ratio of fiber lengths compared to the untreated sample. For analysis, we show a boxplot of the ratio of IdU: CldU measurements (Figure 5). The hydroxyurea-treated sample received hydroxyurea during incorporation of CldU. Therefore the CldU segment of the fiber should be shorter due to the activation of the replication checkpoint by the hydroxyurea induced replication stress. As a result, the ratio of IdU: CldU in the treated sample should be reduced compared to same ratio in the untreated sample. Consistently FiberAI-generated values show that the hydroxyurea-treated sample does have a significantly reduced ratio (pval < 0.001 in 2-sample t test).

**Figure 5:**
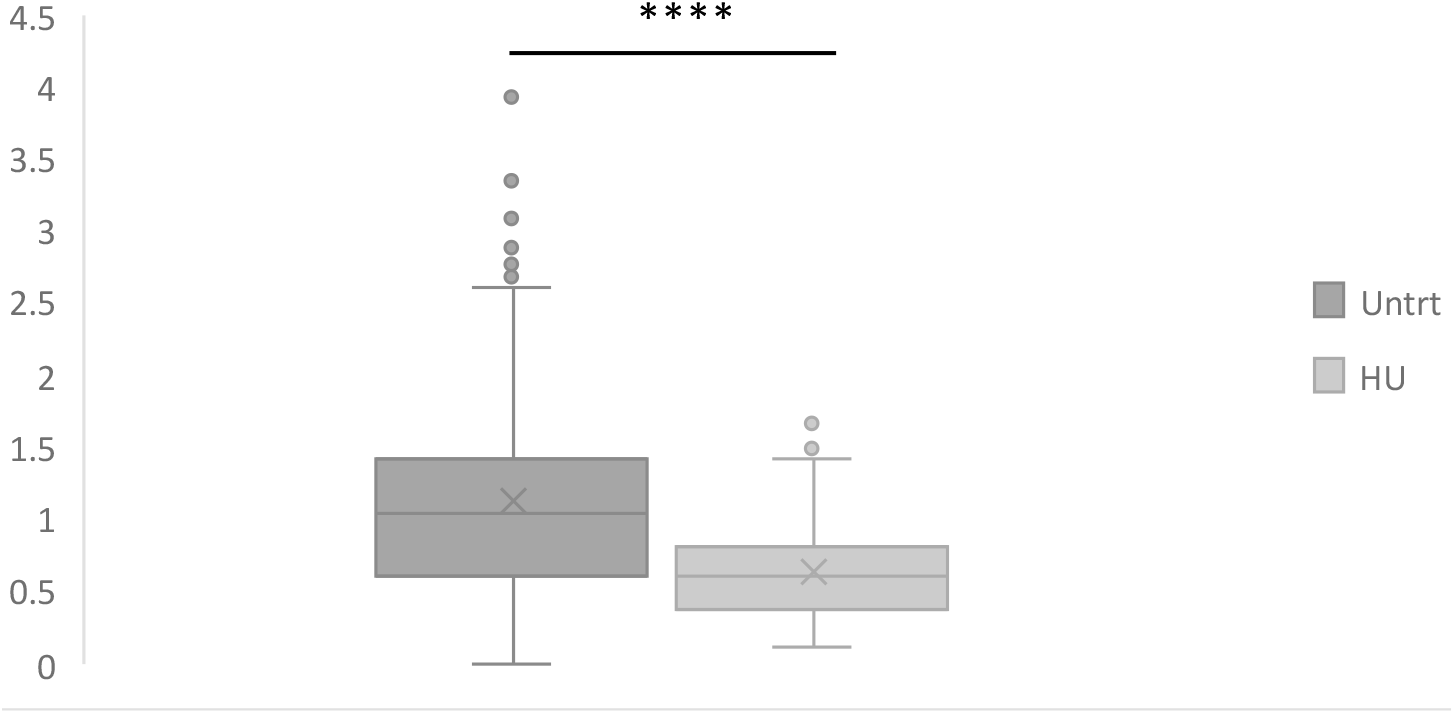
FiberAI-generated ratios (IdU:CldU) for Untreated vs HU-treated samples: FiberAI was used to generate measuremnts for a sample of untreated cells and a sample of cells with incorporation of hydroxyurea (HU) after CldU. FiberAI-generated values shows that the HU-treated sample does have a significantly smaller ratio (pval < 0.001 in 2-sample t test).

## Conclusions

The DNA fiber assay is one of the most powerful and versatile ways to evaluate the integrity and mechanisms that control DNA replication. It provides direct, quantitative information about the progression of the replication fork and a cell’s response to processes that induce replication stress. Analyzing fiber microscopy images can be extremely tedious. There is a lack of free, open-source programs to accurately measure fibers and allow a user to view automated measurements, make their own measurements or insert a manual measurement for any fiber they think the model missed or incorrectly measured. We used deep learning to train an instance segmentation model to find DNA fibers in fiber microscopy images. We provide evidence that our trained model performs very well on object detection and segmentation tasks and post-processing generates highly accurate measurements. We have integrated this model with a graphical interface that allows a user to view automated selections, make their own manual selections, and pick which automated selections they want to keep. Our combined model and graphical interface package is FiberAI, and is provided as free, open source application. With the latest advances in deep-learning, FiberAI is a highly precise and accurate application that solves the tiresome problem of selecting two-component DNA fibers in fiber microscopy images and facilitates high throughput analyses. In this way we expect that FiberAI will contribute in elucidating the molecular processes that ensure the fidelity of DNA replication.

## Data Availability

Data and code available at https://github.com/awezmm/FiberAI

## Author Contributions

Conceptualization: FG and AM. Software: AM, Materials, experiments, and resources: SA, AM, and FG. Writing, Review, and Editing: AM, FG, SA, AK.

